# Cerebral grey matter density is associated with neuroreceptor and neurotransporter availability: A combined PET and MRI study

**DOI:** 10.1101/2020.01.29.924530

**Authors:** Sandra Manninen, Tomi Karjalainen, Lauri J. Tuominen, Jarmo Hietala, Valtteri Kaasinen, Juho Joutsa, Juha Rinne, Lauri Nummenmaa

**Author notes:** Address correspondence to Sandra Manninen, Turku PET Centre and Turku University Hospital, FI-20520 Turku, Finland.

## Abstract

Positron emission tomography (PET) can be used for *in vivo* measurement of specific neuroreceptors and transporters using radioligands, while voxel-based morphometric analysis of magnetic resonance images allows automated estimation of local grey matter densities. However, it is not known how regional neuroreceptor or transporter densities are reflected in grey matter densities. Here, we analyzed brain scans retrospectively from 325 subjects and compared grey matter density estimates with three different neuroreceptors and transporter availabilities. µ-opioid receptors (MORs) were measured with [^11^C]carfentanil (162 scans), dopamine D2 receptors with [^11^C]raclopride (91 scans) and serotonin transporters (SERT) with [^11^C]MADAM (72 scans). The PET data were modelled with simplified reference tissue model. Voxel-wise correlations between binding potential and grey matter density images were computed. Regional binding of all the used radiotracers was associated with grey matter density in region and ligand-specific manner independently of subjects’ age or sex. These data show that grey matter density and MOR and D2R neuroreceptor / SERT availability are correlated, with effect sizes (r^2^) ranging from 0.04 to 0.69. This suggests that future studies comparing PET outcome measure different groups (such as patients and controls) should take grey matter density differences between the groups into account.

## Introduction

Neuroreceptors constitute a major class of targets for pharmacological treatments in psychiatric and neurological conditions. Positron emission tomography (PET) allows *in vivo* measurement of neuroreceptors using radioactive tracer molecules that bind to their target receptors or transporters (1). Accordingly, PET is widely used for quantifying differences in neuroreceptor availabilities across patient and control subjects to investigate pathophysiological changes in specific neurotransmitter circuits (2–4).

However, brain tissue composition related factors associated with radiotracer binding in humans have remained poorly understood. Mesoscopic changes in grey and white matter densities can be derived from magnetic resonance imaging using voxel-based morphometry (VBM) (5). This method is based on segmenting the T1-weighted magnetic resonance (MR) images into grey (GM) and white matter (WM), and comparing the normalized tissue density maps across subject groups. It allows quantification of gross atrophy on the basis of the T1-weighted MR images, yet it yields molecularly unspecific results and cannot pinpoint the neuroreceptor systems involved in the tissue atrophy. The GM signal derived from structural MRI’s is assumed to reflect the gross density of neurons, as a large bulk of cell bodies and neuropils (including glial cells, unmyelinated axons and dendrites) are located in grey matter (6). Consequently, it is possible that binding of PET receptor /transporter radioligands could be associated with the underlying mesoscopic differences in cortical volumes across subjects, as measured with VBM.

In line with this prediction, both molecular and structural neuroimaging studies have found spatially concordant alterations in the brain across multiple conditions. First, GM density declines significantly towards the old age (7,8), and paralleling effects are also observed in opioidergic (9) and dopaminergic (10) neurotransmitter systems, as measured with PET. Similarly, in conditions such as morbid obesity where significant cerebral atrophy is found using VBM (11,12), PET studies show downregulation of opioid (13,14) and possibly also D2-like dopamine receptors (15). Numerous psychiatric conditions including major depression are also associated with significant cerebral atrophy (16) and also regional downregulation of specific neurotransmitters, such as the serotonergic (2,17) and opioidergic (18) systems. Finally, correlational studies linking brain scans with behavior also suggest linkage between cortical volumes and radioligand binding. For example, personality traits associated with pro-sociality are positively associated with both increased frontal cortical volumes (19) and μ-opioid receptor (MOR) availability corresponding regions, as measured with PET (20,21). Few previous studies have already proved a positive correlation with grey matter density and receptor availability with certain receptors (dopamine- and serotonin) (22–24). These studies however investigated single tracers with quite small sample sizes. Here, the focus is to reveal a possible link between grey matter *density* and receptor availability in whole brain area with 3 different tracers to reveal whether tracer uptake and brain density are associated in receptor and transporter specific fashion.

### The current study

The objective of the current study was to investigate the potential links between availability of specific neuroreceptors and transporters measured with PET and cortical density (measured with VBM-and MRI). We focused specifically on µ-opioid and dopamine D2-like receptors and serotonin transporters, as these systems have different and yet partially overlapping distributions in the brain (25–27). We pooled together data from altogether 325 adults who had undergone a PET study with one radioligand each, as well as structural MR imaging with a T1-weighted sequence. We computed voxel-wise as well as region-of-interest based association between GM density and ligand-wise outcome measures. We found that receptor and trasnporter availabilities were positively associated with GM density in regionally specific fashion for each ligand, and these effects were also independent of age and sex related changes.

## Materials and Methods

### Subjects

The data were historical subjects studied at Turku PET Centre. Altogether 325 subjects’ (151 females, age range 18–74, *M*_age_ = 35,8, *SD*_age_ = 13.9 years) scans were included in the study, with a total of 162 [^11^C]carfentanil, 72 [^11^C]MADAM, and 91 [^11^C]raclopride scans (Table 1). The study was conducted in accordance with the Helsinki Declaration. Because the study was a retrospective, register based investigation of historical data, informed consent was waived. The study protocol was reviewed by the institutional review board of the Hospital District of South-Western Finland. The subject-pool consisted primarily (82%) of healthy controls (139 / 162 subjects for carfentanil, 48 / 72 subjects for MADAM and 80 / 91 subjects for raclopride). For the sake of brevity, we included all eligible scans (controls and patients) in the dataset. To rule out potential disease-related confounds in the results, primary analyses were replicated with a control-only sample.

**Table 1.** Subject characteristics for each radioligand used in the study.

| **PET camera** | **n(males)** | **n(females)** | **Age ± SD** | **Dose ± SD** |
| --- | --- | --- | --- | --- |
| **<u>HRRT</u>** |  |  |  |  |
| [<sup>11</sup>C]carfentanil | 39 | 40 | 42±9 | 468±59 |
| [<sup>11</sup>C]MADAM | 32 | 40 | 43±12 | 490±30 |
| [<sup>11</sup>C]raclopride | 35 | 0 | 24±2 | 299±62 |
| **<u>PET-CT</u>** |  |  |  |  |
| [<sup>11</sup>C]carfentanil | 10 | 19 | 36±14 | 252±10 |
| [<sup>11</sup>C]raclopride | 0 | 19 | 43±15 | 253±17 |
| **<u>PET-MRI</u>** |  |  |  |  |
| [<sup>11</sup>C]carfentanil | 54 | 0 | 25±5 | 251±14 |
| **<u>ECAT</u>** |  |  |  |  |
| [<sup>11</sup>C]raclopride | 4 | 11 | 62±10 | 198±15 |
| **<u>HR+</u>** |  |  |  |  |
| [<sup>11</sup>C]raclopride | 0 | 19 | 23±4 | 309±112 |
| **<u>GE-advance</u>** |  |  |  |  |
| [<sup>11</sup>C]raclopride | 0 | 3 | 54±8 | 182±4 |

### MRI and PET data acquisition

Anatomical MR images (voxel size 1 mm^3^) were acquired with Philips Gyroscan Intera 1.5T scanner or Philips Ingenuity 3T PET/MR scanner using T1-weighted sequences. PET data were acquired with the Philips Ingenuity PET/MR scanner, GE Healthcare Discovery 690 PET/CT scanner (General Electric Medical Systems, Milwaukee, WI, USA), GE Advance, CTI-Siemens ECAT EXACT HR+, Siemens HR+ and brain-dedicated high-resolution PET scanner (ECAT HRRT, Siemens Medical Solutions in Turku PET) Centre. MOR availability was measured with [^11^C]carfentanil, D2R availability with [^11^C]raclopride and SERT availability with [^11^C]MADAM.

### MRI Data Preprocessing for Voxel-Based Morphometry

Prior to analysis, the image quality was checked visually and the origo of each T1-weighted image was set to anterior commissure. Structural images were analyzed with SPM12 (www.fil.ion.ucl.ac.uk/spm/) software and the DARTEL pipeline that first creates a study-specific template and for grey and white matter based on the normalized subject-wise input data, and normalizes the subject-wise images using these templates. Default parameter values were used in the DARTEL analysis. The Dartel-normalized GM images were resampled into 2 mm isotropic voxel size and smoothed using a Gaussian kernel of 6 mm full width at half maximum (FWHM).

### PET data preprocessing

To correct for head motion, PET images were first realigned frame-to-frame. Next, they were co-registered with the anatomical MR images. Tracer-wise reference regions (medial occipital cortex for [^11^C]carfentanil and cerebellum for [^11^C]MADAM and [^11^C]raclopride) were automatically defined using FreeSurfer parcellations from the anatomical MR images. The resulting parcellations for occipital and cerebellar grey matter were slightly compressed to avoid spillover from adjacent white matter and cortical regions with specific binding (28). Neuroreceptor/-transporter availability was expressed in terms of *BP*_ND_, which is the ratio of specific to non-displaceable binding in brain. *BP*_ND_ was calculated applying basis function method for each voxel using the simplified reference tissue model (29). *BP*_ND_ is not confounded by differences in peripheral distribution or radiotracer metabolism. For the employed radiotracers, the *BP*_ND_ is traditionally interpreted by target molecule density (Bmax), even though [11C]carfentanil and [11C]raclopride are also sensitive to endogenous neurotransmitter activation. Accordingly, the *BP*_ND_ for these radiotracers should be interpreted as availability of receptors, which in turn is an indirect index of density of available receptors (Bavail).

Resulting subject-wise parametric *BP*_ND_ images were then spatially normalized using the individual flow fields derived from the DARTEL analysis, resampled into 2mm isotropic voxel size to match with the GM segments, and finally smoothed with a Gaussian kernel of 6 mm FWHM. **Figure 1** shows the mean *BP*_ND_ distribution for each radioligand.

**Figure 1.**
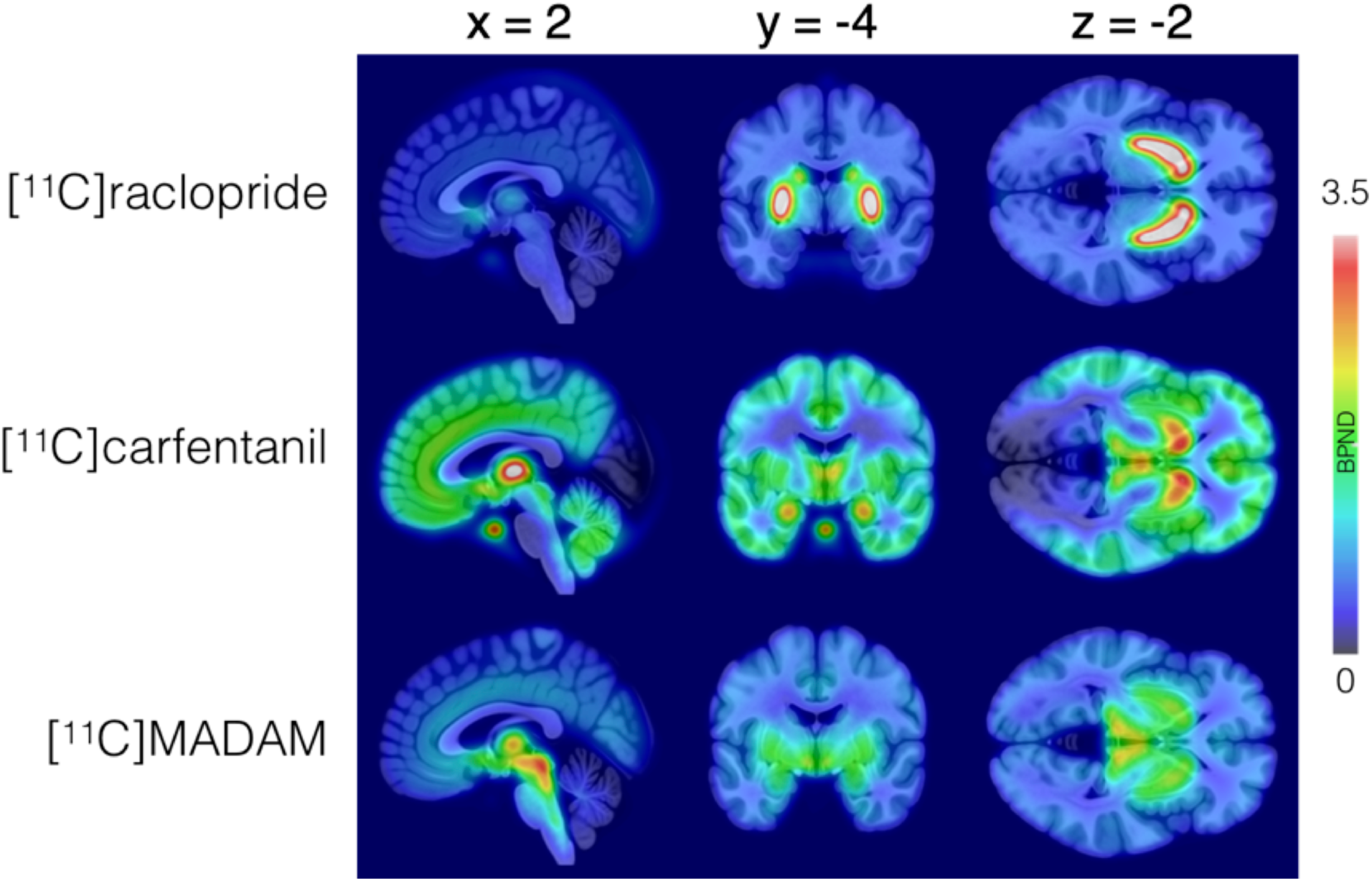
Mean *BP*_ND_-map for each tracer shown over T1-weighted structural MR template image.

### Data analysis

Voxelwise analyses were performed with Matlab R2016b (Math Works, Natick, MA). Regional analyses were performed with R statistical software (https://cran.r-project.org) and *lmer* package. To quantify the association between the tracer-wise *BP*_ND_ and grey matter densities, we computed the voxel-wise correlation between normalized GM segments and each normalized *BP*_ND_ image. Non-parametric Spearman correlation was used in the full-volume analysis to provide additional robustness for the potentially noisy single-voxel data. In the resulting association maps, the voxel intensities reflect the strength of the association (correlation) between the GM/WM density and the *BP*_ND_. To control for false positives, the data were thresholded at p < 0.05, FDR-corrected at cluster level (30). Effects of age and sex were controlled in a separate analysis using general linear model. Because partial volume effects might confound the PET data, we also run a separate analysis for the data where voxelwise partial volume correction was implemented with the PETPVE12 toolbox (31).

Finally, in a complementary methodological approach we also computed the associations between mean GM density and *BP*_ND_‘s in selected a priori regions of interest (the amygdala, caudate, dorsal anterior cingulate cortex (dACC), rostral ACC, thalamus, anterior insula, posterior insula, posterior superior temporal sulcus putamen, nucleus accumbens, precentral gyrus, postcentral gyrus, and orbitofrontal cortex (OFC)). Regions of interests (ROIs) were delineated manually on the mean tracer-specific grey matter DARTEL templates. Finally, to control for the effect of potential confounders for the association between *BP*_ND_ and GMD (age, MRI scanner and PET cameras and their interactions) the ROI analyses were performed using linear mixed models (LMMs).

## Results

We first evaluated the voxel-wise associations between each tracer’s *BP*_ND_ and voxel-wise GM densities. This revealed regionally selective and ligand-specific coupling between *BP*_ND_ and GM density (**Figure 2)**. Please see https://neurovault.org/collections/JOGHJBVA/ for the 3D results images. For [^11^C]raclopride, significant positive associations were found in the putamen and caudate nucleus as well as in thalamus. As expected, no associations were found in areas outside the striatum and thalamus, where [^11^C]raclopride binding is mostly unspecific (26). For [^11^C]carfentanil, positive associations were found in the prefrontal cortex, thalamus, hypothalamus, amygdala, basal ganglia, and cerebellum. With both [^11^C]carfentanil and [^11^C]MADAM, significant negative correlations were found in the putamen, but not in the OFC. For [^11^C]MADAM, positive associations were found in the thalamus, internal capsule, globus pallidus, hippocampal area and parietal cortices. Associations were absent in the reference region (cerebellum), but also remarkably in the frontal cortices where the radioligand does bind to. As expected, the correlations between *BP*_ND_ and GM density were absent in the reference regions for all radio-ligands. The pattern of results remained unchanged even when controlling for voxelwise partial volume effects in the data.

**Figure 2.**
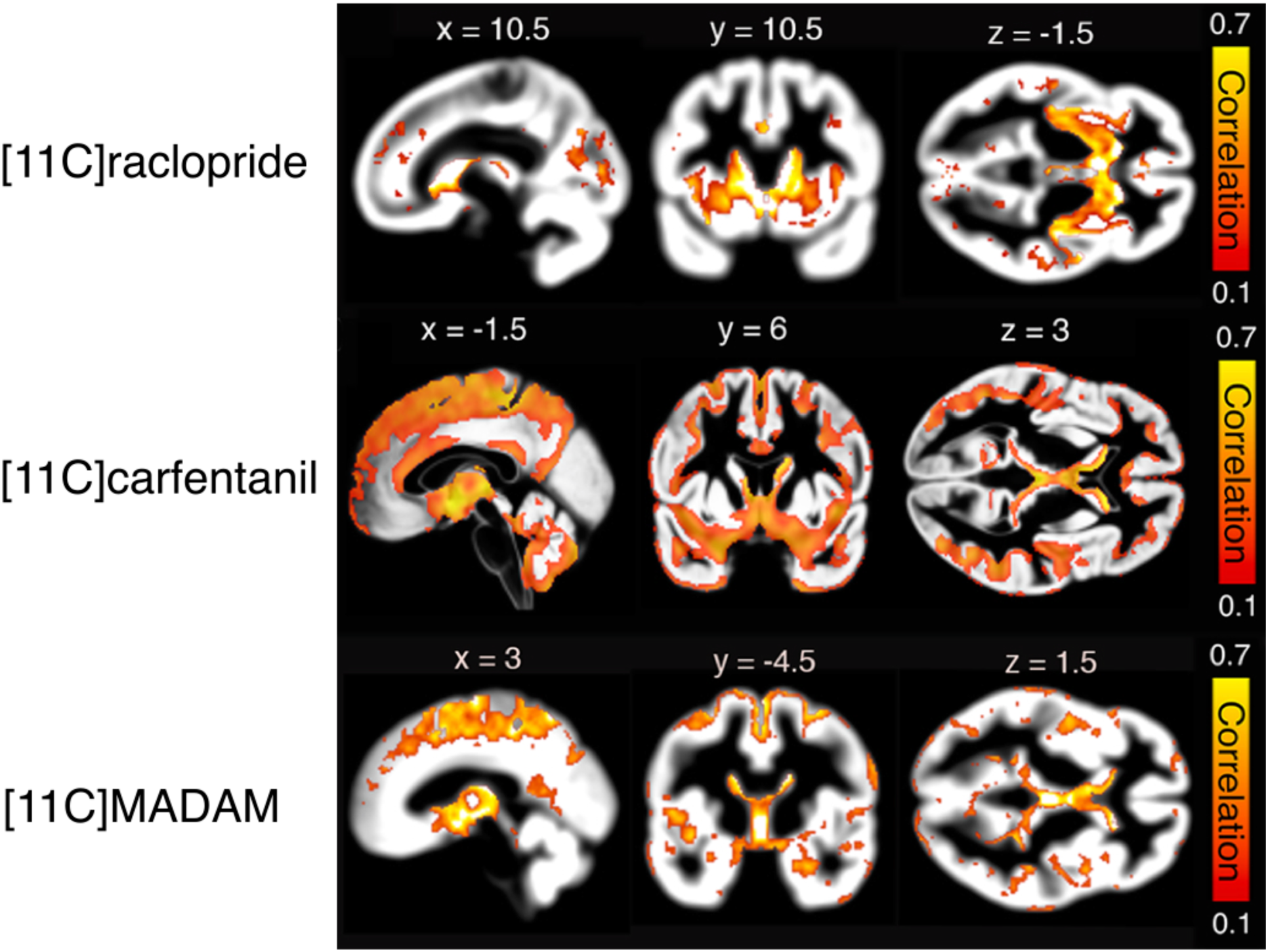
Voxel-wise association maps (Spearman correlation) between grey matter density and *BP*_ND_ for [^11^C]raclopride, [^11^C]carfentanil and [^11^C]MADAM. Colorbar shows correlation coefficient range. All data are thresholded at p < 0.05, FDR corrected at cluster level.

### Control analyses for age, sex, and scanner-related confounds

Ageing is associated with both µ-opioid, dopamine D2 and serotonin transporter availability and GM density. Thus, it is possible that the observed associations between GM and *BP*_ND_ simply reflect age and sex-dependent differences in GM and radiotracer binding, which might explain the positive associations between GM density and *BP*_ND_. To account for this possibility, we also ran a full-volume multiple regression analysis where we predicted the GM density with *BP*_ND_ while controlling with age and sex. Parametric model was deemed appropriate, given that functionally homogenous regions have approximately Gaussian distribution of radioactivity as measured with PET (32). The effects remained essentially unchanged, confirming that the *BP*_ND_-GM-association does not just reflect a confounding effect of age or sex-dependent decline in GM (or *BP*_ND_).

The results from the full-volume analyses were corroborated by the ROI-level statistics (**Figure 3 and Table 2-3**): For [^11^C]raclopride, consistent positive association was found in the striatum and thalamus. For [^11^C]MADAM and [^11^C]carfentanil positive associations were found in the amygdala, thalamus, ventral striatum and caudate nucleus, negative association in the putamen (all p’s < 0.05) and no association in the OFC. Results for the controls-only sample yielded essentially similar results as for the whole sample, the major exception being dampened effects in OFC and putamen for [11C]MADAM. However, even in these regions the correlations were in similar direction as in the full sample, and lack of significance likely arises from compromised statistical power. Finally, because different scanners may yield slightly different binding estimates (despite using *BP*_ND_ as the outcome measure), we also used linear mixed model to predict regional BPND with GMD while accounting for age, MRI scanner, PET camera and their interactions. This revealed that although there were scanner disparities and age dependent effects on *BP*_ND_, these did not account for the association between regional *BP*_ND_ and GMD.

**Table 2.** Bivariate Pearson’s correlations between regional *BP*_ND_ and GM density for each ligand in the whole sample and in healthy controls only. Correlations shown with boldface are significant at p < 0.05. Note: Amygdala and orbitofrontal cortex are not analyzed for [^11^C]raclopride due to lack of specific binding in these regions.

| | [$^{11}C$]carfentanil | | [$^{11}C$]raclopride | | [$^{11}C$] MADAM | |
| --- | --- | --- | --- | --- | --- | --- |
|  | All subjects | Healthy controls | All subjects | Healthy controls | All subjects | Healthy controls |
| Amygdala | **0.24** | **0.25** |  |  | **0.32** | **0.3** |
| Caudate | **0.39** | **0.42** | **0.83** | **0.65** | **0.39** | **0.49** |
| Orbitofrontal cortex | -0.14 | **-0.17** |  |  | **-0.32** | 0.12 |
| Putamen | **-0.32** | **-0.33** | **0.47** | **0.31** | **-0.34** | -0.23 |
| Thalamus | **0.44** | **0.47** | **0.77** | **0.45** | **0.42** | **0.55** |
| Ventral striatum | **0.22** | **0.18** | **0.7** | **0.44** | **0.39** | **0.33** |

**Figure 3.**
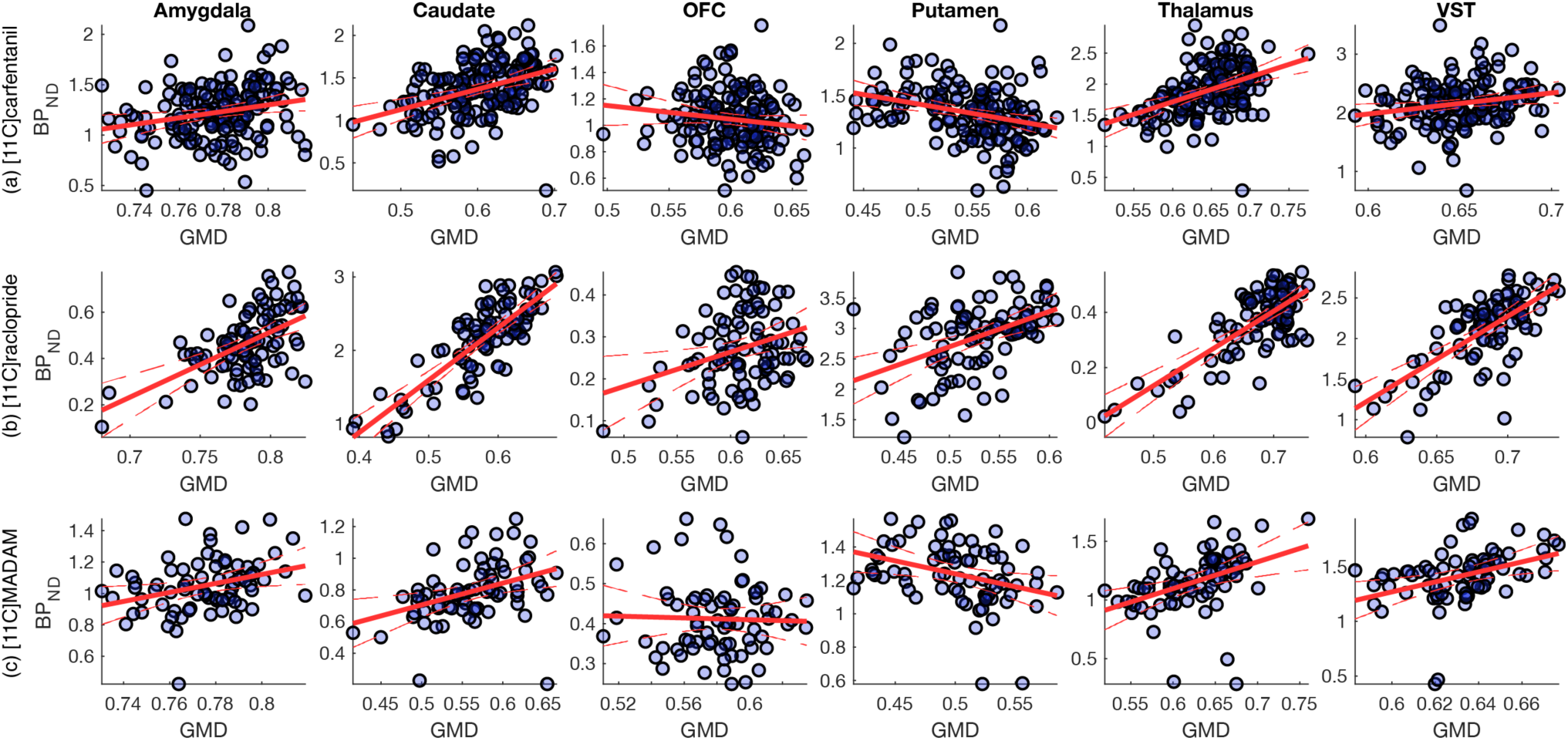
Regional associations and least-squares regression lines between GM density and *BP*_ND_ for all tracers.

**Figure 4.**
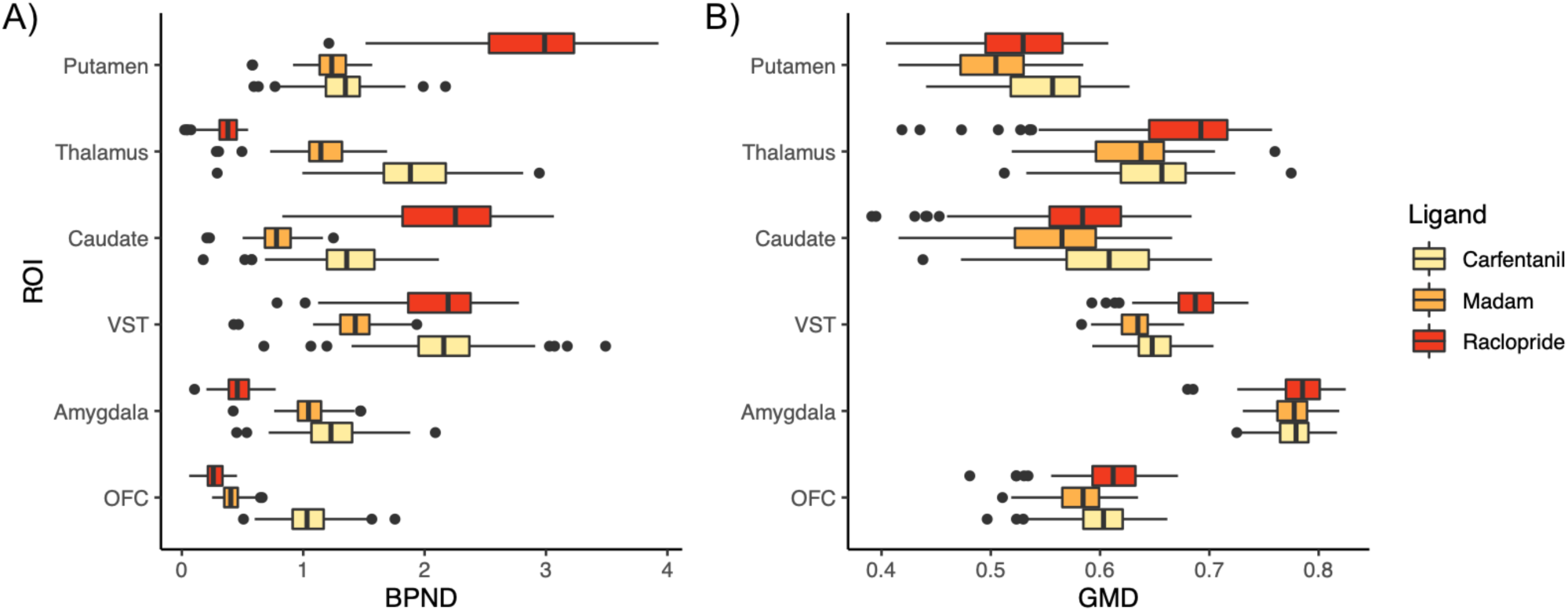
ROI-wise distributions of *BP*_ND_‘s (A) and mean grey matter densities (GMD) for subjects scanned with each tracer. Note: [^11^C]raclopride does not have specific binding outside the striatum, but the mean values are shown for the sake of comparison.

## Discussion

Our main finding was that there is a significant positive association between GM density and *BP*_ND_ with all tested radiotracers. With [^11^C]raclopride the association was strongest in the striatum, and with [^11^C]carfentanil and [^11^C]MADAM in the striatum, thalamus and amygdala. These effects were concordant in the voxel-level and ROI-based analyses. The effects remained essentially unchanged even when controlling for age and sex, which are known to influence both GM and the neuroreceptor systems (7,9). In comparison with previous investigations on GM density and radiotracer uptake, we also observed effects with significantly larger extent, likely due to improved statistical power (22–24). In general, the association between GMD and *BP*_ND_ were of moderate magnitude. Effect sizes (r^2^) of grey matter density on *BP*_ND_ were in the rank of 0.1 for [11C]carfentanil, 0.2 for [11C]raclopride and 0.15 for [11C]MADAM. Although these effects were smaller when controlling for other confounds, they are still considerable given that e.g. 10% between-groups difference in neuroreceptor availability is already substantially large. Altogether these results suggest that availability of these neuroreceptors and transporters, as measured with PET, is reflected in the density of the cerebral tissue as measured with MRI.

### Effects of cortical density on receptor availability

Because MRI-based GM estimate is unspecific with respect to the neuroreceptor and transporter expressions in the neurons in the tissue, we expected to see unique spatial distribution of the GM density and *BP*_ND_ associations for each radiotracer, and not simply significant coupling for all radiotracers in regions with VBM tissue densities. This assumption was confirmed by our data. We observed positive correlations with *BP*_ND_ and GM density with all tracers even though the spatial distribution of these effects varied from tracer to tracer and were specific with respect to the distribution of the targeted system. For example, significant positive association between GM and *BP*_ND_ was found in temporal cortex for [^11^C]carfentanil but not for [^11^C]MADAM. In similar vein, the associations were also found in areas with low radiotracer binding, for example, in cortical midline for [^11^C]MADAM and [^11^C]carfentanil (see **Figure 1** and **Figure 2**). In line with our predictions, mesoscopic variation in cortical density was positively associated with radioligand-specific *BP*_ND_ (that can be presumed to reflect the amount of available receptors in the target tissue). As an important negative control, no associations between *BP*_ND_ and GM density were observed in regions with little or no specific radioligand binding (e.g. frontal cortex for [11C]raclopride).

The most straightforward explanation for the effects is that GM density correlated with receptor / tracer density, because higher grey matter density has larger capacity for receptor / transporter expression. However, one post mortem study found that striatal dopamine transporter availability was not associated with the density of nigral neuron, which project to the striatum (33). Neurotransmission system alterations are typically related to common degenerative brain diseases: dopamine dysfunctions in nigral area and changes in opioid receptor availability are involved in Parkinson’s disease (34,35) and on the other hand, synaptic loss and serotonergic dysfunction correlates with severity of the symptoms in Alzheimer’s disease (1,36). Accordingly, it is possible that a functional neurotransmission is dependent on available and functioning receptors on the neuron cell surface and not solely the exact amount of the neurons. Furthermore, all neurons don’t necessarily express the same amount of different transmitters’ receptors in central nervous system. However, we also unexpectedly observed negative associations between radioligand binding and GM density in the putamen for [^11^C]carfentail and [^11^C]MADAM. We have no clear explanation for this effect, but it has been previously speculated that such effects may reflect regional differences in the regional neurotransporter / receptor availabilities, or the consistency of the segmentation of the T1-weightened images (22–24). Or, when the regional GM density grows, there might be relatively more such neurons, that don’t express those neurotransmitter’s receptors in question.

Finally, in numerous regions (such as caudate) we observed a positive association with GM density and all *BP*_ND_ of all the radiotracers. This is not unexpected, as VBM-based grey matter density indices the gross number of cell bodies and neuropil (Purves et. al 2018), and single GM voxel may contain tens of thousands of glial cells and neurons with different receptor expression profiles. It is well known that the VBM-based density metrics are difficult to interpret on their own (37) and the present data shows that intersubject variability in neuroreceptor and transporter availability may influence these MR-driven metrics. In any case, from methodological perspective these finding suggest that mere (unspecific) grey matter differences might confound concomitant neuro-receptor-level changes between, for example, patient and control groups. Hence, it would be advisable to statistically test the unique effects of group membership and underlying cortical atrophy in neuro-receptor PET studies. Altogether these results thus suggest that *BP*_ND_ and GM densities are coupled: If someone has high grey matter tissue density, he/she also likely has high tracer binding potentials (expect for in putamen for raclopride and MADAM). However, this association can be manifested in both regions having high and low *BP*_ND_ or GM density, thus it not simply driven by high-binding areas in PET or high-density areas in in MR. We also stress that this coupling is far from complete, as the correlations between BP_ND_ and GM densities vary from 0.22 to 0.83. The proportion of GMD not accounted a radiotracer’s BP_ND_ likely reflects both error variance and presence of neurons operating with other neurotransmitters within each GMD voxel, yet these effects cannot be partialed out in the current design where each subject was scanned with a single radioligand.

### Effects of age on brain morphology and neuroreceptor level changes

Atrophy and synaptic loss are consistently observed in healthy aging brain (Gunning-Dixon et al., 2010; Peelle et al. 2012; Resnick et al. 2003), but also contributing to degenerative late age pathologies such as Alzheimer’s and Parkinson’s diseases (36,40–43). Similarly, specific neurotransmitter systems show age-dependent alterations. While dopamine D2 (44,45) and serotonin transporter (46) densities decrease during aging, an opposite pattern is observed for µ-opioid receptors (47). Analysis of the associations between ligand-specific BP_ND_ and GM density revealed strong regional association that were, to some extent, modulated by age and sex. However, the overall pattern of results (i.e. association between BP_ND_:s and GM densities) remained essentially unchanged even when controlling for age and sex, thus the association cannot be explained by age-dependent atrophy that would lower radiotracer binding in general. It must be noted that our subjects’ were primarily young and middle-aged adults (mean age 35.8 years), and effects of age-dependent atrophy on radiotracer uptake might be observed only in significantly older subjects. Nevertheless the fusion analysis suggests that joint analysis of cerebral atrophy (VBM) and receptor/transporter systems could be useful in tracking the development of neurological and psychiatric conditions: Regional volumetric brain changes are associated with specific neuropsychological symptoms, and also to specific neurotransmitter system alterations. The presently observed linkage between transporter and receptor densities highlights that it is imperative to investigate how the linkage is manifested across neurological and psychiatric pathologies.

## Limitations

We combined retrospective data from several different studies performed with different scanners. However, calibrated PET scanners yield consistent data (48) and the employed outcome measure (*BP*_ND_) should theoretically control for minor differences in scanner signal-to-noise ratios (29). In line with this, controlling for scanner type did not in general alter the findings (Table 2). Despite this, even *BP*_ND_ estimates are known to vary between-scanner differences may be amplified depending on implemented PVEc (49). Thus, caution should be warranted when combining data across PET cameras, and the dependencies between camera types and independent variables (such as age, sex, and patient status) should be taken into account and optimally controlled for when building up the pooled sample. The scans for different radioligands came from different subjects, so individual differences in the tracer-specific samples might also confound the results. The direct causal linkage between the neuroreceptor and transporter availability and GM density cannot be established using the current cross-sectional data, thus we do not know if increase in the number of density of specific neuroreceptors or transporters would yield increase in the GM signal in MR. Future translational and animal imaging studies are needed for elucidating this aspect. Finally, future studies could address the relationship between receptor and transporter density and other MRI-derived metrics, such as cortical density estimated with surface analyses.

## Conclusions

We conclude that regional binding of [^11^C]carfentanil, [^11^C]raclopride and [^11^C]MADAM is associated with cortical density at the binding site. Grey matter density and tracer *BP*_ND_ are associated in region and ligand-specific manner for [^11^C]carfentanil, [^11^C]raclopride and [^11^C]MADAM. These data thus suggest that VBM signal is coupled on specific neuroreceptor and transporter levels, and also suggests that when comparing any PET outcome measure across two groups, it is advisable to consider GM density differences between the group as well.

## Conflict of Interest

The authors declare no conflict of interest

## Acknowledgments

This work was supported by the Academy of Finland (grants #304385 and #294897) and Sigrid Juselius Foundation grants to LN, and the Alfred Kordelin Foundation grant to SM, and Päivikki and Sakari Sohlberg Foundation and Finnish Cultural Foundation Varsinais-Suomi Regional Fund to TK.

## Data availability

Per current Finnish legislation and research permission the dataset cannot be publicly shared.

## Author contributions

Designed the study: TK, LJT, JR, LN

Acquired data: SM, LJT, JH, VK, JJ, JR

Analyzed the data: SM, TK

Wrote the manuscript: SM, TK, LJT, JH, VK, JJ, JR, LN

---

**Table 2.** Results of linear mixed models where regional ligand-specific BP_ND_ were predicted with age, GMD, MRI and PET scanners and their interactions when applicable. Effects whose SE:s do not overlap zero are shown in boldface. Note: Amygdala and orbitofrontal cortex are not analyzed for [^11^C]raclopride due to lack of specific binding in these regions.

| | | [$^{11}C$]carfentanil | | | [$^{11}C$]raclopride | | | [$^{11}C$]MADAM | | |
| --- | --- | --- | --- | --- | --- | --- | --- | --- | --- | --- |
|  |  | Estimate | SE | T | Estimate | SE | T | Estimate | SE | T |
| Amygdala | Age | -0.09 | 0.02 | **-4.32** |  |  |  | -0.01 | 0.02 | -0.60 |
|  | GMD | 0.04 | 0.02 | **2.02** |  |  |  | 0.05 | 0.02 | **2.47** |
|  | MRI scanner 1 | 0.19 | 0.04 | **5.10** |  |  |  |  |  |  |
|  | MRI scanner 2 |  |  |  |  |  |  |  |  |  |
|  | Age X GMD | 0.00 | 0.01 | 0.27 |  |  |  | -0.01 | 0.02 | -0.39 |
|  | GMD X MRI scanner 1 | 0.00 | 0.03 | 0.03 |  |  |  |  |  |  |
|  | GMD X MRI scanner 2 |  |  |  |  |  |  |  |  |  |
| Caudate | Age | 0.07 | 0.03 | **2.69** | -0.09 | 0.03 | **-3.30** | 0.00 | 0.03 | -0.04 |
|  | GMD | 0.12 | 0.03 | **4.35** | 0.18 | 0.03 | **5.92** | 0.09 | 0.03 | **2.78** |
|  | MRI scanner 1 | 0.22 | 0.05 | **4.07** | 0.16 | 0.05 | **3.47** |  |  |  |
|  | MRI scanner 2 |  |  |  | 0.07 | 0.13 | **0.51** |  |  |  |
|  | Age X GMD | -0.02 | 0.02 | -0.73 | -0.02 | 0.02 | -1.27 | 0.02 | 0.03 | 0.48 |
|  | GMD X MRI scanner 1 | 0.04 | 0.04 | 0.81 | -0.10 | 0.04 | **-2.47** |  |  |  |
|  | GMD X MRI scanner 2 |  |  |  | -0.02 | 0.05 | -0.30 |  |  |  |
| OFC | Age | 0.06 | 0.02 | **2.39** |  |  |  | 0.02 | 0.03 | 0.55 |
|  | GMD | -0.04 | 0.02 | **-1.70** |  |  |  | 0.00 | 0.03 | -0.04 |
|  | MRI scanner 1 | 0.16 | 0.04 | **4.42** |  |  |  |  |  |  |
|  | MRI scanner 2 |  |  |  |  |  |  |  |  |  |
|  | Age X GMD | 0.03 | 0.01 | **2.04** |  |  |  | -0.03 | 0.02 | -1.12 |
|  | GMD X MRI scanner 1 | 0.03 | 0.03 | 0.90 |  |  |  |  |  |  |
|  | GMD X MRI scanner 2 |  |  |  |  |  |  |  |  |  |
| Putamen | Age | 0.04 | 0.02 | **2.03** | -0.08 | 0.03 | **-2.52** | 0.00 | 0.02 | 0.06 |
|  | GMD | 0.02 | 0.04 | 0.45 | 0.09 | 0.04 | **2.26** | 0.05 | 0.02 | **-1.98** |
|  | MRI scanner 1 | 0.22 | 0.05 | **4.43** | 0.08 | 0.06 | **1.43** |  |  |  |
|  | MRI scanner 2 |  |  |  | -0.11 | 0.13 | -0.87 |  |  |  |
|  | Age X GMD | 0.02 | 0.02 | 0.76 | 0.02 | 0.03 | 0.89 | 0.01 | 0.03 | -0.21 |
|  | GMD X MRI scanner 1 | 0.00 | 0.05 | 0.08 | -0.04 | 0.04 | -0.93 |  |  |  |
|  | GMD X MRI scanner 2 | - | - | - | -0.13 | 0.08 | **-1.66** |  |  |  |
| Thalamus | Age | -0.05 | 0.02 | **-2.16** | -0.09 | 0.06 | **-1.55** | 0.00 | 0.04 | -0.10 |
|  | GMD | 0.04 | 0.02 | **1.82** | 0.29 | 0.07 | **4.40** | 0.08 | 0.03 | **2.24** |
|  | MRI scanner 1 | 0.11 | 0.04 | **2.56** | 0.12 | 0.09 | **1.38** |  |  |  |
|  | MRI scanner 2 |  |  |  | -0.15 | 0.21 | -0.74 |  |  |  |
|  | Age X GMD | 0.00 | 0.02 | 0.24 | 0.06 | 0.04 | **1.78** | -0.03 | 0.04 | -0.74 |
|  | GMD X MRI scanner 1 | 0.05 | 0.03 | **1.36** | -0.17 | 0.08 | **-2.05** |  |  |  |
|  | GMD X MRI scanner 2 |  |  |  | -0.32 | 0.09 | **-3.57** |  |  |  |
| VS | Age | -0.03 | 0.02 | **-1.66** | -0.05 | 0.03 | **-1.37** | 0.01 | 0.03 | 0.36 |
|  | GMD | 0.04 | 0.02 | **2.23** | 0.10 | 0.03 | **3.16** | 0.08 | 0.03 | **2.99** |
|  | MRI scanner 1 | 0.19 | 0.04 | **4.93** | 0.02 | 0.05 | 0.48 |  |  |  |
|  | MRI scanner 2 |  |  |  | -0.31 | 0.14 | **-2.20** |  |  |  |
|  | Age X GMD | 0.00 | 0.02 | -0.12 | 0.01 | 0.02 | 0.42 | 0.02 | 0.02 | -0.80 |
|  | GMD X MRI scanner 1 | 0.01 | 0.03 | 0.20 | -0.04 | 0.04 | -0.84 |  |  |  |
|  | GMD X MRI scanner 2 |  |  |  | -0.18 | 0.06 | **-2.95** |  |  |  |

**Table S-1**. Original studies whose data was selected for the analysis. Note: Not all scans from all projects are included.

## Notes

https://neurovault.org/collections/JOGHJBVA/

## REFEERENCES

1. Heiss W-D, Herholz K. Brain Receptor Imaging*. J Nucl Med. 2006;47(2):302–12.

2. Gryglewski G, Lanzenberger R, Kranz GS, Cumming P. Meta-analysis of molecular imaging of serotonin transporters in major depression. J Cereb Blood Flow Metab. 2014;34(7):1096–103.

3. Volkow ND, Fowler JS, Wang GJ, Baler R, Telang F. Imaging dopamine’s role in drug abuse and addiction. Neuropharmacology. 2009;56(SUPPL. 1):3–8.

4. Whone AL, Watts RL, Stoessl AJ, Davis M, Reske S, Nahmias C, et al. Slower progression of Parkinson’s disease with ropinirole versus levodopa: The REAL-PET study. Ann Neurol. 2003;54(1):93–101.

5. Ashburner J, Friston KJ. Voxel-Based Morphometry - The Methods. Neuroimage. 2000;11(1):805–21.

6. Purves D et al. NEUROSCIENCE. OXFORD UNIV PRESS; 2018.

7. Resnick SM, Pham DL, Kraut MA, Zonderman AB, Davatzikos C. Longitudinal Magnetic Resonance Imaging Studies of Older Adults : A Shrinking Brain. J Neurosci. 2003;23(8):3295–301.

8. Tang Y, Whitman GT, Lopez I, Baloh RW. Brain volume changes on longitudinal magnetic resonance imaging in normal older people. J Neuroimaging. 2001;11(4):393–400.

9. Zubieta J-K, Dannals RF, Frost JJ. Gender and Age Influences on Human Brain μ-Opioid Receptor Binding Measured by PET. Am J Psychiatry. 1999;156(6):842–8.

10. Morgan DG. The dopamine and serotonin systems during aging in human and rodent brain. A brief review. Prog Neuropsychopharmacol Biol Psychiatry. 1987;11:153–7.

11. Karlsson HK, Tuulari JJ, Hirvonen J, Lepomäki V, Parkkola R, Hiltunen J, et al. Obesity is associated with white matter atrophy: A combined diffusion tensor imaging and voxel-based morphometric study. Obesity. 2013;21(12):2530–7.

12. Tuulari JJ, Karlsson HK, Antikainen O, Hirvonen J, Pham T, Salminen P, et al. Bariatric Surgery Induces White and Grey Matter Density Recovery in the Morbidly Obese: A Voxel-Based Morphometric Study. Hum Brain Mapp. 2016;37(11):3745–56.

13. Karlsson HK, Tuulari JJ, Tuominen L, Hirvonen J, Honka H, Parkkola R, et al. Weight loss after bariatric surgery normalizes brain opioid receptors in morbid obesity. Mol Psychiatry. 2016 Aug;21(8):1057–62.

14. Karlsson HK, Tuominen L, Tuulari JJ, Hirvonen J, Parkkola R, Helin S, et al. Obesity Is Associated with Decreased -Opioid But Unaltered Dopamine D2 Receptor Availability in the Brain. J Neurosci. 2015;35(9):3959–65.

15. Wang GJ, Volkow ND, Logan J, Pappas NR, Wong CT, Zhu W, et al. Brain dopamine and obesity. Lancet. 2001;357(9253):354–7.

16. Goodkind M.. Identification of a Common Neurobiological Substrate for Mental Illness. 2015;72(4):305–315.

17. Spies M, Knudsen GM, Lanzenberger R, Kasper S. The serotonin transporter in psychiatric disorders: Insights from PET imaging. The Lancet Psychiatry. 2015;2(8):743–55.

18. Kennedy SE, Koeppe R a, Young E a, Zubieta J-K. Dysregulation of endogenous opioid emotion regulation circuitry in major depression in women. Arch Gen Psychiatry. 2006;63(11):1199–208.

19. Lewis PA, Rezaie R, Brown R, Roberts N, Dunbar RIM. Ventromedial prefrontal volume predicts understanding of others and social network size. Neuroimage. 2011;57(4):1624–9.

20. Manninen S, Tuominen L, Dunbar RI, Karjalainen T, Hirvonen J, Arponen E, et al. Social Laughter Triggers Endogenous Opioid Release in Humans. J Neurosci. 2017;37(25):6125–31.

21. Nummenmaa L, Manninen S, Tuominen L, Hirvonen J, Kalliokoski KK, Nuutila P, et al. Adult attachment style is associated with cerebral u-opioid receptor availability in humans. Hum Brain Mapp. 2015;36(9):3621–8.

22. Kraus C, Hahn A, Savli M, Kranz GS, Baldinger P, Höflich A, et al. Serotonin-1A receptor binding is positively associated with gray matter volume - A multimodal neuroimaging study combining PET and structural MRI. Neuroimage. 2012;63(3):1091–8.

23. Woodward ND, Zald DH, Ding Z, Riccardi P, Ansari MS, Baldwin M, et al. Cerebral morphology and dopamine D2/D 3receptor distribution in humans: A combined [18 F]fallypride and voxel-based morphometry study. 2010;46(1):31–8.

24. Winkler et al. Cortical thickness or Grey Matter volume. 2011;53(3):1135–46.

25. Frost JJ, Wagner HNJ, Dannals RF, Ravert HT, Links JM, Wilson AA, et al. Imaging opiate receptors in the human brain by positron tomography. J Comput Assist Tomogr. 1985;9(2):231–6.

26. Egerton A, Mehta MA, Montgomery AJ, Lappin JM, Oliver D. The dopaminergic basis of human behaviors : a review of molecular imaging studies. Neurosci Biobehav Rev. 2013;33(7):1109–32.

27. Chalon S, Tarkiainen J, Garreau L, Hall H, Emond P, Vercouillie J, et al. 4-methylphenyl thio) benzylamine as a Ligand of the Serotonin Transporter with High Affinity and Selectivity. Pharmacology. 2003;304(1):81–7.

28. Karjalainen, T., Santavirta, S., Kantonen, T., Tuisku, J., Tuominen, L., Hirvonen, J., Hietala, J., Rine, J., Nummenmaa L. Magia: Robust automated modeling and image processing toolbox for PET neuroinformatics. Frontiers in Neuroinformatics. Front Neuroinform. 2020;

29. Gunn RN, Lammertsma AA, Hume SP, Cunningham VJ. Parametric Imaging of Ligand-Receptor Binding in PET Using a Simplified Reference Region Model. Neuroimage [Internet]. 1997 Nov 1 [cited 2018 Nov 22];6(4):279–87. Available from: https://www.sciencedirect.com/science/article/pii/S1053811997903037?via%3Dihub

30. Benjamini Y, Hochberg Y. Benjamini Y, Hochberg Y. Controlling the false discovery rate: a practical and powerful approach to multiple testing. J R Stat Soc B. 1995;57(1):289–300.

31. Gonzalez-Escamilla G, Lange C, Teipel S, Buchert R, Grothe MJ. PETPVE12: an SPM toolbox for Partial Volume Effects correction in brain PET – Application to amyloid imaging with AV45-PET. Neuroimage [Internet]. 2017;147:669–77. Available from: http://www.sciencedirect.com/science/article/pii/S1053811916308023

32. Teymurazyan A, Riauka T, Jans H-S, Robinson D. Properties of Noise in Positron Emission Tomography Images Reconstructed with Filtered-Backprojection and Row-Action Maximum Likelihood Algorithm. J Digit Imaging [Internet]. 2013;26(3):447–56. Available from: https://doi.org/10.1007/s10278-012-9511-5

33. Saari L, Kivinen K, Gardberg M, Joutsa J, Noponen T, Kaasinen V. Dopamine transporter imaging does not predict the number of nigral neurons in Parkinson disease. Neurology. 2017 Mar;

34. Heiss W-D, Hilker R. The sensitivity of 18-fluorodopa positron emission tomography and magnetic resonance imaging in Parkinson’s disease. Eur J Neurol. 2004 Jan;11(1):5–12.

35. Piccini P, Weeks RA, Brooks DJ. Alterations in opioid receptor binding in Parkinson’s disease patients with levodopa-induced dyskinesias. Ann Neurol. 1997;42(5):720–6.

36. Terry RD. Cell death or synaptic loss in Alzheimer disease. J Neuropathol Exp Neurol. 2000;59(12):1118–9.

37. Zatorre RJ, Fields RD, Johansen-Berg H. Europe PMC Funders Group Plasticity in Gray and White : Neuroimaging changes in brain structure during learning. 2013;15(4):528–36.

38. Peelle JE, Cusack R, Henson RNA. Adjusting for global effects in voxel-based morphometry: Gray matter decline in normal aging. Neuroimage. 2012;60(2):1503–16.

39. Gunning-Dixon FM, Brickman AM, Cheng JC, Alexopoulos GS. Aging of Cerebral White Matter: A Review of MRI Findings. Int J Geriatr Psychiatry. 2010;24(2):109–17.

40. Alves J, Soares JM, Sampaio A, Gonçalves ÓF. Posterior cortical atrophy and Alzheimer’s disease: A meta-analytic review of neuropsychological and brain morphometry studies. Brain Imaging Behav. 2013;7(3):353–61.

41. Benson D, Davis R, BD S. Posterior cortical atrophy. Arch Neurol. 1988 Jul;45(7):789–93.

42. Tuite PJ, Mangia S, Michaeli S. Magnetic Resonance Imaging (MRI) in Parkinson’s Disease. J Alzheimer’s Dis Park. 2013;Suppl 1:1.

43. Bellucci A, Mercuri NB, Venneri A, Faustini G, Longhena F, Pizzi M, et al. Parkinson’s disease: From synaptic loss to connectome dysfunction. Neuropathol Appl Neurobiol. 2016;42(1):77–94.

44. Rinne JO, Hietala J, Ruotsalainen U, Säkö E, Laihinen A, Någren K, et al. Decrease in Human Striatal Dopamine D 2 Receptor Density with Age: A PET Study with [^11^ C]Raclopride. J Cereb Blood Flow Metab. 1993;13(2):310–4.

45. Antonini A, Leenders KL, Reist H, Thomann R, Beer HF, Locher J. Effect of age on D2 dopamine receptors in normal human brain measured by positron emission tomography and 11C-raclopride. Arch Neurol. 1993 May;50(5):474–80.

46. van Dyck CH, Malison RT, Seibyl JP, Laruelle M, Klumpp H, Zoghbi SS, et al. Age-related decline in central serotonin transporter availability with [(123)I]beta-CIT SPECT. Neurobiol Aging. 2000;21(4):497–501.

47. Zubieta JK, Dannals RF, Frost JJ. Gender and age influences on human brain mu-opioid receptor binding measured by PET. Am J Psychiat. 1999;156(June):842–8.

48. Johansson J, Teuho J, Linden J, Tuna U, Tolvanen T, Saunavaara V, et al. Image quantification in high-resolution PET assessed with a new anthropomorphic brain panthom. IEEE Nuclear Science Symposium and Medical Imaging Conference (2013 NSS/MIC). 2013;(IEEE Conference):1–7.

49. Schain M, Tóth M, Cselényi Z, Stenkrona P, Halldin C, Farde L, et al. Quantification of serotonin transporter availability with [11C]MADAM — A comparison between the ECAT HRRT and HR systems. Neuroimage [Internet]. 2012;60(1):800–7. Available from: http://www.sciencedirect.com/science/article/pii/S1053811911014492

## [11C]carfentanili scans

Viikari J, Raitakari O, Keltikangas-Järvinen L, Hietala J (2012) Temperament trait Harm Avoidance associates with µ-opioid receptor availability in frontal cortex: a PET study using [(11)C]carfentanil. Neuroimage 61, 670–676.

Tuulari, J.J., Tuominen, L., de Boer, F., Hirvonen, J., Nuutila, P., & Nummenmaa, L. (2017). Feeding releases endogenous opioids in humans. The Journal of Neuroscience, 37, 8284–8291.

Saanijoki, T., Tuominen, L.J., Nummenmaa, L., Arponen, E., Kalliokoski, K., & Hirvonen, J. (2018). Opioid release after high-intensity interval training in healthy human subjects. Neuropsychopharmacology, 43, 246–254.

Manninen, S., Tuominen, L., Dunbar, R.I.M., Karjalainen, T., Hirvonen, J., Arponen, E., Hari, R., Jääskeläinen, I.P., Sams, M., & Nummenmaa, L. (2017). Social laughter triggers endogenous opioid release in humans. The Journal of Neuroscience, 37, 6125–6131;

Nummenmaa, L., Tuominen, L., Dunbar, R., Hirvonen, J., Manninen, S., Arponen, E., Machin, A., Hari, R., Jääskeläinen, I.P., & Sams, M. (2016). Social touch modulates endogenous µ-opioid system activity in humans. NeuroImage, 138, 242–247.

Karlsson, H.K., Tuominen, L., Tuulari, J.J., Hirvonen, J., Parkkola, R., Helin, S., Salminen, P., Nuutila, P., & Nummenmaa, L. (2015). Obesity is associated with decreased mu-opioid but unaltered dopamine D2 receptor availability in the brain. The Journal of Neuroscience, 35, 3959–3965.

## [11C]MADAM scans

Tuominen, L., Salo, J., Hirvonen, J., Naåren, K., Laine, P., Melartin, T., Isometsä, E., Viikari, J., Cloninger, C.R., Raitakari, O., Hietala, J., & Keltikangas-Järvinen, L. (2012) Temperament, character and serotonin activity in the human brain: a positron emission tomography study based on a general population cohort. Psychological Medicine, 43, 881–894.

Majuri, J., Joutsa, J., Johansson, J., Voon, V., Alakurtti, K., Parkkola, R., … Kaasinen, V. (2016). Dopamine and Opioid Neurotransmission in Behavioral Addictions: A Comparative PET Study in Pathological Gambling and Binge Eating. Neuropsychopharmacology, 42, 1169–1177

## [11C]raclopride scans

Aalto, S., Ingman, K., Alakurtti, K., Kaasinen, V., Virkkala, J., Någren, K., Rinne, J.O., Scheinin H. (2015). Intravenous ethanol increases dopamine release in the ventral striatum in humans: PET study using bolus-plus-infusion administration of [(11)C]raclopride. Journal of Cerebral Blood Flow and Metabolism, 35, 424–31.

Alakurtti, K., Aalto, S., Johansson, J.J., Någren, K., Tuokkola, T., Oikonen, V., Laine, M., & Rinne J.O. (2011). Reproducibility of striatal and thalamic dopamine D2 receptor binding using [11C]raclopride with high-resolution positron emission tomography. Journal of Cerebral Blood Flow and Metabolism, 3, 555–565.

Bäckman, L., Waris, O., Johansson, J., Andersson, M., Rinne, J.O., Alakurtti, K., Soveri, A., Laine, M., & Nyberg, L. (2017). Increased dopamine release after working-memory updating training: Neurochemical correlates of transfer. Scientific Reports, 7(1):7160.

Kemppainen, N., Ruottinen, H., Någren, K., & Rinne, J.O. (2000). PET shows that striatal dopamine receptors are differentially affected in AD. Neurology, 55, 205–209.

Rinne, J.O., Laihinen, A., Någren, K., Bergman, J., Solin, O., Haaparanta, M., Ruotsalainen, U., Rinne, U.K. (1990) PET demonstrates different behaviour of striatal dopamine D-1 and D-2 receptors in early Parkinson’s disease. Journal of Neuroscience. Research, 27, 494–499.

